# Optogenetic confirmation of transverse-tubular membrane excitability in intact cardiac myocytes

**DOI:** 10.1101/2023.11.29.569144

**Authors:** M. Scardigli, M. Pásek, L. Santini, C. Palandri, E. Conti, C. Crocini, M. Campione, L. Loew, A. A. F. de Vries, D. A. Pijnappels, F. S. Pavone, C. Poggesi, E. Cerbai, R. Coppini, P. Kohl, C. Ferrantini, L. Sacconi

## Abstract

T-tubules (TT) form a complex network of sarcolemmal membrane invaginations, essential for well-coordinated excitation-contraction coupling (ECC) and, thus, homogeneous mechanical activation of cardiomyocytes. ECC is initiated by rapid depolarization of the sarcolemmal membrane. Whether TT membrane depolarisation is active (local generation of action potentials; AP) or passive (following depolarisation of the outer cell surface sarcolemma; SS) has not been experimentally assessed in cardiomyocytes. Based on the assessment of ion flux pathways needed for AP generation, we hypothesise that TT are excitable. We therefore explored TT excitability experimentally, using an all-optical approach to stimulate and record trans-membrane potential changes in TT that were electrically insulated from the SS membrane by transient osmotic shock. Our results establish that cardiomyocyte TT can generate AP. These AP show electrical features that differ substantially from those observed in SS, consistent with differences in the density of ion channels and transporters in the two different membrane domains. We propose that TT-generated AP represent a safety mechanism for TT AP propagation and ECC, which may be particularly relevant in pathophysiological settings where morpho-functional changes reduce the electrical connectivity between SS and TT membranes.

**KEY POINTS:**

- Cardiomyocytes are characterized by a complex network of membrane invaginations (the T-tubular system) that propagate action potentials to the core of the cell, ensuring synchronous and uniform cell contraction.
- In this study, we investigate whether the T-tubular system is able to generate action potentials autonomously, rather than following depolarization of the outer cell surface sarcolemma.
- For this purpose, we developed a fully optical platform to probe and manipulate the electrical dynamics of sub-cellular membrane domains.
- Our findings demonstrate that T-tubules are intrinsically excitable, revealing distinct characteristics of self-generated T-tubular action potentials.
- This active electrical capability may serve as a protective mechanism against voltage drops occurring within the T-tubular network.

## INTRODUCION

Mammalian cardiomyocytes contain a complex network of sarcolemmal membrane invaginations referred to as the transverse/axial tubular system, or the T-tubule (TT) system. In healthy cardiomyocytes, TT membranes are continuous with the surface sarcolemma (SS). SS and TT jointly mediate coordinated electrical and mechanical activation of individual cardiomyocytes (Tidball *et al*., 1991) by supporting excitation-contraction coupling (ECC) across the entire cell (Franzini-Armstrong *et al*., 1973; Bers, 2002).

Despite indications of non-homogeneous distribution of ion flux pathways and chemical gradients in SS and TT compartments, there are no significant differences between action potential (AP) profiles recorded in SS and TT membranes of healthy cardiomyocytes, presumably because the two membrane compartments are electrically well-coupled (Sacconi *et al*., 2012). The characteristic length constant of electrotonic potential decay in TT is one order of magnitude larger than a typical cardiomyocyte radius (Scardigli *et al*., 2017), indicating a remarkable safety factor for coordinated voltage changes across the entire sarcolemma. This may be impaired, however, in cardiac diseases where substantial TT remodelling has been observed (Sacconi *et al*., 2012; Crocini *et al*., 2014).

This poses an interesting, and as yet experimentally unresolved question: is TT membrane depolarisation, and hence well-coordinated initiation of ECC across the entire cell volume, dependent on active AP generation in TT, or driven by passive electrotonic effects from the SS membrane?

We therefore implemented an experimental investigation which involved, first, electrical insulation of TT from SS membranes and, second, an all-optical approach to locally depolarise TT or SS membranes while simultaneously tracking changes in their trans-membrane potential. Electrical insulation of TT from SS membranes was achieved by transient exposure of cells to an acute osmotic shock (an approach called ‘acute detubulation (Kawai *et al*., 1999; Ferrantini *et al*., 2014), which ties-off TT near their junctions with the SS membrane). Using random-access multi-photon (RAMP) microscopy and cardiomyocytes containing a combination of either CheRiff or ChR2 (optogenetic actuators) (Hochbaum *et al*., 2014) and di-4-AN(F)EPPTEA (voltage-sensitive dye; VSD) (Yan *et al*., 2012), we found that TT membranes are electrically excitable, i.e. able to generate local AP even after electrical insulation from the SS.

## METHODS

### Animals

We used (i) 12-week-old female transgenic mice (ChR2-mhc6-cre +) with cardiomyocyte-specific expression of ChR2 (H134R variant) and (ii) age-matched mice from of the same strain (C57BL/6J). Animal handling and procedures were performed in accordance with guidelines in Directive 2010/63/EU of the European Parliament (protection of animals used for scientific purposes). The experimental protocol was approved by the Italian Ministry of Health (authorization number 944/2018-PR).

### Adeno-associated virus-based gene transfer of CheRiff

Mouse hearts were driven to express the optogenetic actuator CheRiff (Hochbaum *et al*., 2014) coupled to enhanced green fluorescent protein (eGFP) by adeno-associated virus (AAV) based transduction using AAV2/9.45.GgTnnt2.CheRiff∼eGFP.WHVPRE.SV40pA particles. This AAV (Pulicherla *et al*., 2011) contains a transgene consisting of the chicken cardiac troponin T promoter, the coding sequence of the blue light-activatable cationic channelrhodopsin CheRiff tagged at its C terminus with eGFP, the woodchuck hepatitis virus posttranscriptional regulatory element, and the simian virus 40 polyadenylation signal. Production, purification and quantification of these vector particles was done as described in Nyns *et al*.(Nyns *et al*., 2017) Systemic delivery of AAV particles was performed as in (Yan *et al*., 2012). Briefly, AAV particles were injected into the left jugular vein of mice anesthetized with isoflurane (3% for induction, 1.5% for maintenance, in oxygen). Body temperature was maintained at 37°C using a homeothermic blanket system (50300; Stoelting Co., USA) and mice were monitored using respiratory rate and toe pinch response tests throughout the procedure. Mice were placed on a stereotaxic apparatus and shaved over the chest to allow identification of the jugular vein underneath the mouse skin, after which the shaved area was sterilized with betadine and 70% ethanol. Next, lidocaine solution was applied to the skin surface and a 5-mm-long cervical incision was made to expose the jugular vein. An insulin syringe (30G×1/6’’, Biotekne, Italy) was used to slowly inject 200 μL of the injection mix composed of ∼2×10^11^ genome copies of AAV diluted in phosphate-buffered saline (P4417, Merck) containing 2.5 mM MgCl_2_ and 1 mM KCl. After stopping of the bleeding with surgical swabs, the skin was sutured with Vicryl Plus Ethicon (VCP394H, Johnson & Johnson Medical N.V., Belgium) and mice were placed back into their cages. Tail vein injection was also explored. To this end, tails were cleaned and sterilized with 70% ethanol and a bent needle of an insulin syringe was inserted into one of the lateral veins, followed by the slow administration of 200 μL of the injection mix. At 4 weeks post-transduction, hearts were excised and used for the isolation of ventricular cardiomyocytes as described below.

### Cardiomyocyte isolation

Left ventricular cardiomyocytes from all experimental groups were isolated by enzymatic dissociation, as described before (Crocini *et al*., 2016b). In brief, animals were heparinized (5,000 U/kg body weight, i.p.) and deeply anaesthetized with isoflurane. The heart was excised, quickly bathed in cell isolation solution at 37 °C, and the ascending aorta was cannulated for perfusion in Langendorff mode. Cell isolation solution is calcium-free and contains (in mM): 113 NaCl, 4.7 KCl, 0.6 KH_2_PO_4_, 0.6 Na_2_HPO_4_, 1.2 MgSO_4_, 12 NaHCO_3_, 10 KHCO_3_, 10 HEPES, 30 taurine, 10 glucose, 10 2,3-butanedionemonoxime, pH 7.3 adjusted with NaOH). After perfusion for 3-4 min with a constant flow of 7-8 mL/min at 37 °C, the cell isolation solution was switched to a recirculating enzyme-containing solution, made from the cell isolation solution by adding 0.1 mg/mL Liberase TM (Merk Life Science S.r.l., Germany). After 7 min, the left ventricle was harvested, cut into small pieces in cell isolation solution supplemented with 1 mg/mL bovine serum albumin, and gently stirred to facilitate dissociation of cardiomyocytes. The cell suspension was allowed to settle at room temperature (RT, 20 °C) for 10 min, before the cell pellet was re-suspended in calcium-free Tyrode buffer solution. Calcium-free Tyrode buffer solution contained (in mM): 133 NaCl, 4.8 KCl, 1.2 MgCl_2_, 10 glucose, 10 μM blebbistatin, 4 μM cytochalasin D and 10 HEPES, pH 7.35 adjusted with NaOH, throughout all experiments. Isolated ventricular cardiomyocytes were gradually re-adapted at RT to calcium by adding 50 or 100 μM CaCl_2_ every 5-8 min, until a concentration of 1 mM CaCl_2_ was reached.

### Sarcolemma labelling

Cells were placed in Tyrode buffer containing 0.5 mM CaCl_2_. Cardiomyocyte staining was performed by adding 2 μg/mL of the VSD di-4-AN(F)EPPTEA (Yan *et al*., 2012) (from 2 mg/mL stock, dissolved in ethanol). After 5 min, cells were resuspended in fresh Tyrode buffer containing 1 mM CaCl_2_. Labelled cardiomyocytes were used for experiments within 30 min. Staining and imaging were performed at RT.

### Acute detubulation

Disconnection of TT membranes from the SS (henceforth called ‘detubulation’) was achieved by hyper-osmotic shock, as previously described (Kawai *et al*., 1999; Ferrantini *et al*., 2014). Briefly, 1.5 M formamide was added to the cell suspension (in Tyrode buffer containing 0.5 mM CaCl_2_). After 15 min, cells were resuspended rapidly in fresh Tyrode buffer containing 1 mM CaCl_2_. The detubulation procedure was performed at room temperature. For RAMP measurements detubulation was induced after sarcolemmal staining with VSD.

### Quantification of T-tubular density

Isolated cardiomyocytes, both intact (CTRL) and formamide-treated (DETUB), were incubated for 20 minutes with 5 µM of Di-3-ANEPPDHQ (Thermo Fisher Scientific, USA) membrane dye. This approach allowed us to investigate the fraction of detached T-tubules because only those still continues with the surface were labelled. CRTL and DETUB cells were resuspended in calcium-free Tyrode buffer, placed into a glass-bottomed chamber (P50G-0-14-F, MatTek, USA) and observed using a 63× oil immersion objective on a Leica TCS5 inverted microscope (Leica, Germany). The excitation wavelength was 488 nm, and the detection window was set to 504-649 nm. A single image was recorded from each cardiomyocyte, choosing a z-plane in the central portion of the cell. TT quantitative analysis was performed using AutoTT (Guo & Song, 2014) on confocal images, as previously described (Crocini *et al*., 2016b). In brief, AutoTT skeletonizes the global architecture of the TT system to extract morphological patterns, discriminating transverse and axial elements of the system. To determine the densities of transverse and axial elements, the total number of pixels in each of these TT sub-categories was divided by the total number of pixels of the intracellular region imaged.

### Patch-clamp experiments

To perform patch-clamp recordings, viable ventricular cardiomyocytes were superfused with Tyrode buffer containing 1.8 mM CaCl_2_ at RT. Patch-clamp data recording and analysis were performed using a Multiclamp700B amplifier in conjunction with pClamp10.0 and a DigiData 1440A AD/DA interface (Molecular Devices, USA). AP recordings and K^+^ current measurements were conducted in whole-cell ruptured patch mode. The pipette solution contained (in mM): 130 K-aspartate, 0.1 Na-GTP, 5 Na_2_-ATP, 11 EGTA, 5 CaCl_2_, 2 MgCl_2_, 10 HEPES, pH 7.2 adjusted with KOH; free [Ca^2+^] was calculated using maxchelator (https://somapp.ucdmc.ucdavis.edu/pharmacology/bers/maxchelator/) and was equal to 243 nM. For K^+^ current measurements, 0.3 mM CdCl_2_ was added to the bath solution. AP trains were electrically elicited with short depolarizing stimuli (3 ms square current pulses, 500-1,000 pA) at 1 Hz pacing rate. Recordings of global potassium current (*I*_K_) were carried out using depolarization steps of 900 ms, from -50 mV to 0 mV, immediately after eliciting the Na current. I_K-peak_ was calculated as the maximum current, while I_K-end_ is the residual outward current at 900 ms. For L-type calcium (*I*_CaL_) and sodium (*I*_Na_) current recordings, the pipette solution contained (in mM) 120 CsCl, 10 HEPES, 5 Na_2_ATP, 10 TEACl, 5 MgCl_2_, 4.3 CaCl_2_ and 10 EGTA (calculated free [Ca^2+^] = 226 nM), pH 7.2 adjusted with CsOH, while the external solution contained (in mM) 135 NaCl, 5.4 CsCl, 10 glucose, 10 HEPES, 1 MgCl_2_, 1.8 CaCl_2_, 0.33 NaH_2_PO_4_, 0.3 BaCl_2_, pH 7.4 adjusted with CsOH. Peak sodium current (*I*_Na_) was elicited with a (15 ms) depolarizing step from -80 mV (holding potential) to -45 mV, while *I*_CaL_ was elicited with a longer (200 ms) depolarization step from -45 mV to 0 mV right immediately eliciting the Na current. *I*_Na_ and *I*_CaL_ were induced at a repeat rate of 1 Hz.

### Modelling

The model used in the present study is based on a previously published model of rat ventricular cardiomyocytes incorporating TT (Pasek *et al*., 2006). The model was modified to reproduce the key properties (e.g. resting membrane potential, AP amplitude and duration, AP rate adaptation, intracellular Ca^2+^ transient amplitude and duration) of the mouse ventricular cardiomyocytes used in this study (Bondarenko *et al*., 2004). Modifications included (i) adjustment of geometric parameters of the model cell; (ii) reformulation of ion flux pathways in SS and TT membranes; (iii) adaptation of sarcoplasmic reticulum Ca^2+^ pump activity; (iv) adjustment of ion exchange between the TT lumen and the extracellular space. All modifications are described in detail in the Supplementary material (Paragraph 1). The model was implemented in MATLAB v.7.2 (MathWorks, Inc.). Numerical computation of the system of 47 nonlinear differential equations was performed using the solver for stiff system ode15s (MATLAB). Model equations were solved simultaneously, using a variable time-step (0.4×10^-6^ - 0.1 s) adjusted to keep the estimated relative error of inner variables below a threshold value of 10^-6^. Steady state behaviour of the model was achieved by running it for 600 s of equivalent cell lifetime at the specified stimulation frequencies. The MATLAB code of the model is available at https://www.it.cas.cz/en/d3/l033.

### Random-access multiphoton microscope and optogenetics stimulation

The basic design of our RAMP microscope has been described elsewhere (Sacconi *et al*., 2012; Crocini *et al*., 2014; Scardigli *et al*., 2017). In the present study, the system was implemented with a fibre-based optical stimulation system for optogenetic pacing. A fibre-coupled LED (M470F3, Thorlabs, Germany) operating at 470 ± 25 nm was employed as fibre-coupled light source, and a 3D micromanipulator was used to place the distal end of the multi-mode optical fibre (FG200UEA, Thorlabs, Germany) in proximity of the cell. The numerical aperture (N.A.) of the optical fibre and the angle of the fibre with respect to the main optical axis were chosen to avoid direct collection of blue light by the RAMP excitation/detection objective (40×, water immersion, N.A. 0.8; Olympus, Japan). This is crucial, given that blue light can generate unwanted phosphorescent signals inside the objective glass that may affect optical recordings (not shown). The RAMP fluorescence signal was detected using a photon-counting module, based on the GaAsP photomultiplier tube (PMT) with an ultra-fast gate circuit (H11706P-40, Hamamatsu, Japan). An emission filter of 655 ± 20 nm was used for fluorescence detection. A trigger distributor switched off the high-voltage power supply circuit of the PMT during optical stimulation. This avoids the possibility that endogenous or exogenous fluorescence signals overload the PMT during blue light illumination. Cells were optically stimulated using pulses of blue light with 3 ms duration and a light intensity of 10 mW mm^-2^ at the output of the optical fibre. Alternatively, cells could be electrically field-stimulated using two parallel platinum wires (⌀ 250 μm), placed in the bath at a distance of 6.3 mm. Square pulses of 10–20 V, lasting 3 ms, were used to trigger AP. In a typical RAMP measurement, we performed one line-scan along the SS, and two to four line-scans on different TT membrane segments (for 20 subsequent trials). The length of scanned lines ranged from 5 to 10 μm, with an integration time per membrane pass of ∼200 μs, leading to a temporal resolution of 0.4 - 2 ms. Opto-electronic components of the setup were computer-controlled with custom-made software developed in LabVIEW 7.1 (National Instruments). Optical recording were analysed with a custom-made software written in LabVIEW 2013 (National Instruments).

### Statistics

For each experimental condition, data from one cell were averaged, and this average was used for comparison and statistical analysis. To assess whether measured data came from a normal, log-normal, or other distribution, we performed a Shapiro-Wilk test of all data before and after their logarithmic transformation. Median was used as the best representation of central data values. In patch-clamp experiments, one-way analysis of variance (ANOVA) tests were used to compare electrophysiological features between CTRL and DETUB. In optical recordings, two-way ANOVA tests were used to compare electrophysiological features between SS and TT at different stimulation modality, the Bonferroni post hoc analysis was used. AP optical stimulation efficiency in CTRL and DETUB was compared using the Fisher exact test. Statistical analysis was performed using OriginPro 2018 (OriginLab).

## RESULTS

### Ion flux pathways across sarcolemma in mouse cardiomyocyte and in silico assessment of TT excitability

Structural and patch-clamp data from ventricular cardiomyocytes which underwent acute detubulation through osmotic shock (DETUB) were compared to intact cells (CTRL) to asses AP-relevant ion current distributions between TT and SS (Figure 1). Our data indicate that L-type Ca^2+^ channels are expressed predominantly in the TT membrane (note: *I*_CaL_ density was reduced by about one half in DETUB, compared to CTRL cells), whereas Na^+^ and K^+^ channels densities appear to be uniform across the whole cell membrane. Consistent with the high *I*_CaL_ density in TT, a reduction of AP duration at 50% and 90% repolarisation was found in DETUB cells, compared to CTRL, while no significant differences in AP amplitude or resting membrane potential were detected between the groups.

**Figure 1:**
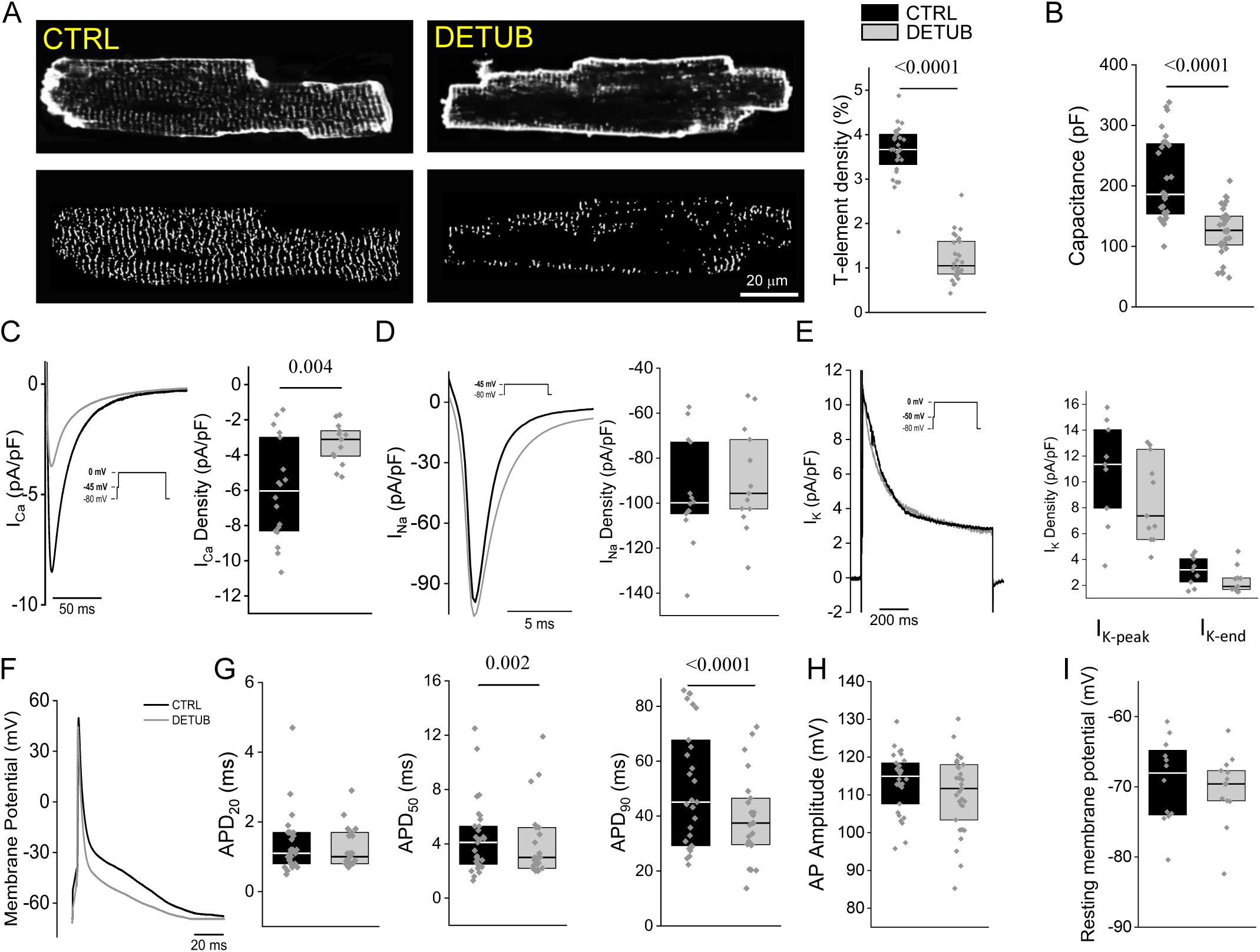
Patch-clamp recordings on CTRL and DETUB cells. (A) Two representative confocal images of intact (CTRL) and detubulated (DETUB) cardiomyocytes. Sarcolemma stained with Di-3-ANEPPDHQ. In the bottom panels, correspondent images of same cells showing skeletonized TT system, obtained with AUTO-TT. On the right, TT density expressed as percentage (%) in CTRL (black box; N = 3 animal, n = 28 cells) and DETUB (grey box; N = 3, n = 32) cardiomyocytes. (B) Cell capacitance (pF), measured by dividing membrane time constant by membrane resistance, recorded in CTRL (N = 6, n = 30) and DETUB (N = 6, n = 31) cardiomyocytes. (C) Representative traces and average values of L-Type Ca^2+^ current (*I*_CaL_) density at 0 mV from CTRL (N = 5, n = 18) and DETUB (N = 5, n = 13) cardiomyocytes. (D) Representative traces and average values of peak Na^+^ current (*I*_Na-peak_) density at - 0 mV from CTRL (N = 5, n = 14) and DETUB (N = 5, n = 13). (E) Representative traces and average values of was calculated as the peak outward K^+^ current (I_K-peak_) and residual outward current at 900 ms (I_Kend_) at 0 mV from CTRL (N=5, n=9) and DETUB (N=5, n=12). (F) Representative AP recorded in CTRL and in DETUB cardiomyocytes, paced at 1 Hz. (G) Duration at 20%, 50% and 90% of repolarization from AP peak (APD_20_, APD_50_, APD_90_, respectively) recorded in CTRL (N = 6, n = 30) and DETUB (N = 6, n = 31) cardiomyocytes, stimulated at 1 Hz. (H) AP amplitude expressed as the difference between the resting membrane potential (RMP) and the maximum potential recorded, stimulated at 1 Hz. (I) RMP. Single data are reported as grey points and box plots with a range of 25^th^ to 75^th^ percentile are superimposed. Median values are represented by the line in box plots. P-values were calculated with ANOVA one-way test using Bonferroni test. N indicates the number of animals and n the number of cells in each data series. Raw data are available at: https://doi.org/10.6084/m9.figshare.23742231.

Measured ion flux pathways were incorporated into an adapted two-compartment computer model of mouse ventricular cardiomyocytes (see supplementary material, paragraph 1). The model predicts that TT are potentially excitable (Figure 2A) and that the AP duration of isolated TT should be longer than in SS alone. Furthermore, the model showed that coordinated ECC could be ensured both with active AP generation, or passive electrotonic conduction, in TT (Figure 2B). It is not possible, therefore, to distinguish between these two possible mechanisms, based on spatially resolved data on ECC and contraction dynamics.

**Figure 2:**
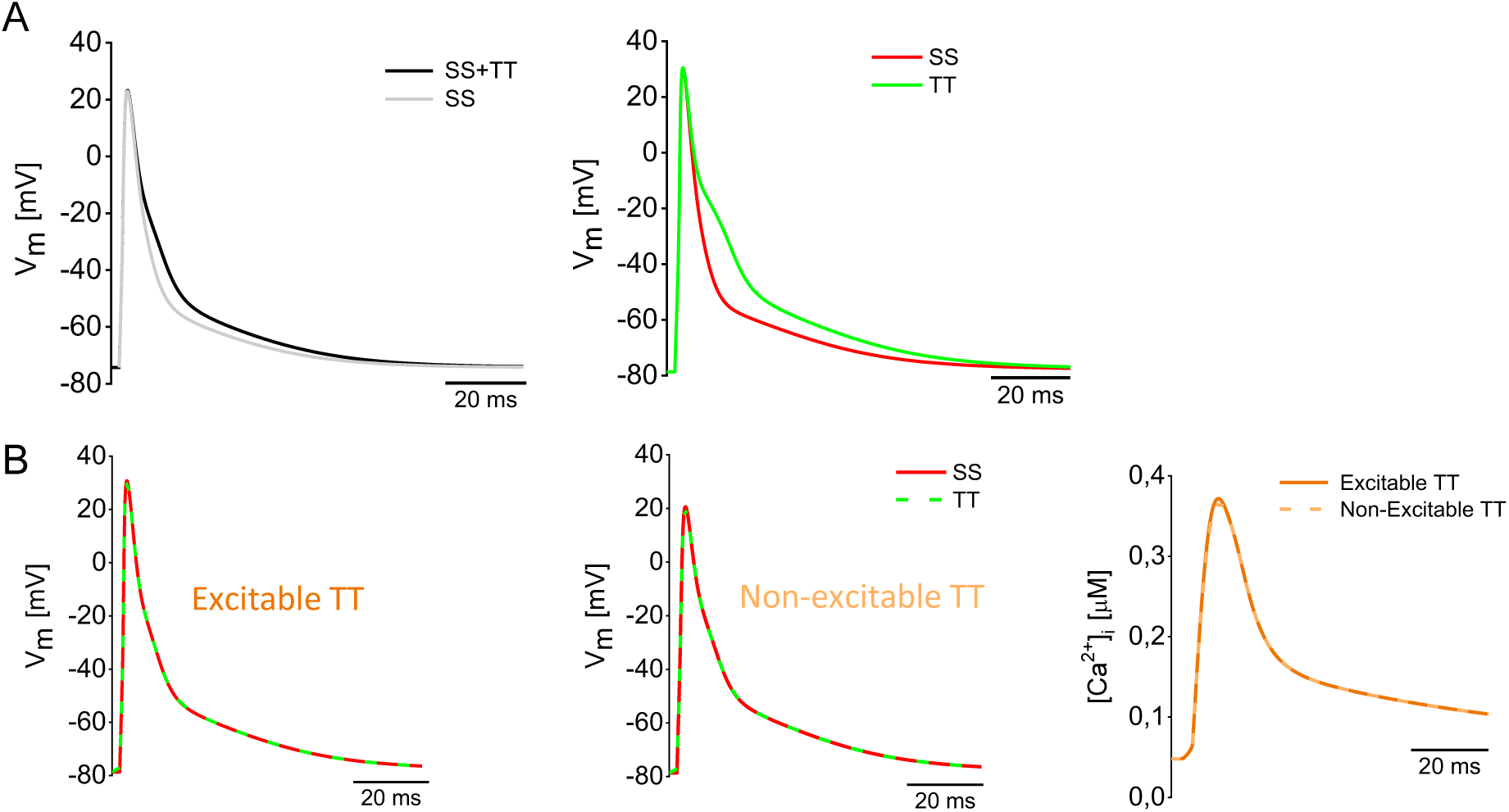
*In silico* prediction of TT membrane excitability. (A) On the left: Simulation of 1 Hz steady-state AP shapes in CTRL (black) and DETUB cells (grey) assuming a residual TT fraction equal to zero. Intracellular ion concentrations were fixed at experimentally clamped values: [Na^+^]_i_ = 10 mM, [K^+^]_i_ = 130 mM and [Ca^2+^]_i_ = 200 nM. Extracellular/tubular ion concentrations were: [Na^+^]_e_ = 140 mM, [K^+^]_e_ = 5.4 mM and [Ca^2+^]_e_ = 1.8 mM. On the right: Simulation of TT (green trace) and SS (red trace) AP after separation of TT. TT lumen ion concentrations were close to the extracellular ion concentration: [Na^+^]_TT_ =140 mM, [K^+^]_TT_ = 5.4 mM and [Ca^2+^]_TT_ = 1.8 mM. (B) Simulated AP shapes in the presence (Excitable TT) and absence (Non-excitable TT) of the fast sodium current in TT (dashed green). Simulated cytosolic Ca^2+^-transient in the presence of excitable TT (dark orange) and non-excitable TT (dashed light orange). Raw data are available at: https://doi.org/10.6084/m9.figshare.23742231.

### Optical induction and recording of AP in SS and TT of CTRL cells

RAMP microscopy (Sacconi *et al*., 2012; Crocini *et al*., 2014) was combined with optogenetic stimulation, employing an ultra-fast gated detection system that was turned off during light stimulation (Figure 3A). This approach was first validated in intact ventricular myocytes, constitutively expressing a light-gated ion channel (ChR2) and labelled with di-4-AN(F)EPPTEA VSD. RAMP rapidly scanned linear segments on SS and TT membranes during electrical or whole-cell optogenetic stimulation. Both stimulation modes allowed AP induction in SS and TT membranes of CTRL cells (Figures 3B,C). As expected, electrically and optically stimulated AP amplitudes and kinetics in TT are not significantly different from those in SS (Figure 3D-F).

**Figure 3:**
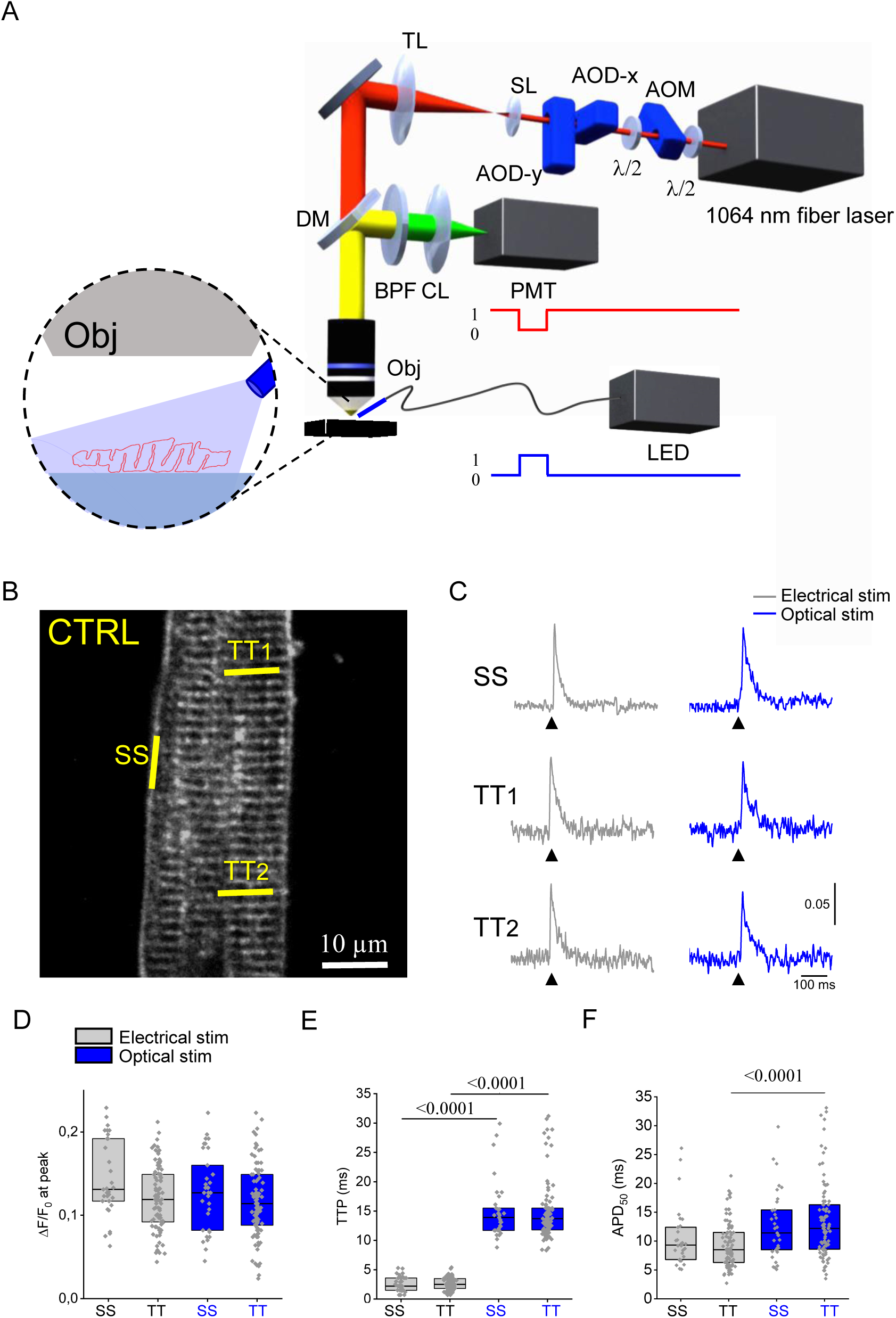
All-optical AP recording in TT und SS membrane sites of CTRL cardiomyocytes. (A) Scheme of the random access multiphoton (RAMP) microscope: 1064-nm fibre laser, an acousto-optic modulator (AOM), two orthogonally mounted acousto-optic deflectors (AOD-x and AOD-y), scanning lens(SL), tube lens (TL), dichroic mirror (DM), water immersion objective (Obj), band-pass filter (BPF), condenser lens (CL), and photo-multiplier tube (PMT). Cells were optically stimulated using a fibre-coupled LED, placing the distal end of the multi-mode optical fibre in the proximity of the cell (highlighted in the inset). A trigger distributor was used to switch-off the PMT (red traces) during optical stimulation (blue trace). (B) Two-photon fluorescence image of a ventricular myocyte constitutively expressing ChR2 and labelled with VSD. The yellow lines mark the probed membrane sites (sarcolemma: SS; T-tubules: TT_1_ and TT_2_). Scale bar 10 μm. (C) Normalized fluorescence traces (ΔF/F_0_) recorded from the three scanned membrane domains (average of 10 sequential runs). An AP (elicited at 200 ms, black arrowheads) is clearly visible in all three sites, regardless of whether the cell was stimulated electrically or optically. AP elicited using electrical field are represented in grey, while APs evoked using optical stimulation are in blue. (D) AP amplitude (ΔF/F_0_ at peak), (E) time-to-peak (TTP) and (F) APD_50_, measured upon electrical (black) or optical (blue) stimulation. Single cell data are reported as grey points and box plots with a range of 25^th^ to 75^th^ percentile are superimposed. Median values are represented by the line in box plots. Data from 33 SS and from 93 TT recordings (N = 6 animals, n = 33 cells). P values were calculated with two-way ANOVA with Bonferroni test. Raw data are available at: https://doi.org/10.6084/m9.figshare.23742231.

The system records a relative fluorescence change (ΔF/F_0_) which, at the AP peak, is in the order of 10 - 15% (Figure 3D). While this provides a sufficient signal-to-noise ratio for assessing AP dynamics, it is lower than what was found in our previous investigation (:: 20 %) (Crocini *et al*., 2016b). This is not unexpected, considering that in the present mouse line (Crocini *et al*., 2016a) the 1064 nm light excites both the VSD and ChR2-conjugated td-Tomato fluorescent protein (Drobizhev *et al*., 2011). In a subset of experiments where cardiomyocytes from CTRL mice were transfected with light-activated channels fused with eGFP (not excited at 1,064 nm), we found (Figure S4) that the relative fluorescence change at the AP peak can be restored to previously reported values, both upon electrical and optical stimulation.

Comparing AP features between the two stimulation modalities, we found a substantially longer time-to-peak AP (roughly 5-fold, 15 ms instead of 3 ms; TTP in Figure 3E) during optical compared to electrical stimulation, as previously reported (Bruegmann *et al*., 2010). Effects of ChR2 activation on AP repolarization (e.g. APD_50_ in Figure 3F) were less pronounced or absent. These results indicate that a full-optical approach can be used to quantitatively assess AP dynamics in different sarcolemmal domains.

### Optical induction and recording of AP in SS and TT of DETUB cells

Next, we employed the same approach on ChR2-expressing cardiomyocytes that underwent acute detubulation. As indicated in Figure 4A, cells were first stained with the VSD and then acutely detubulated by osmotic shock. Thereafter, AP were optically recorded from SS and TT membranes upon electrical or optical stimulation (Figure 4B). As expected, upon electrical stimulation, TT that were detached from the SS did not exhibit AP (grey traces), showing no voltage membrane variation above the shot-noise (:: 5 mV). This means that structurally detached and electrically uncoupled TT were not excited by either the local extracellular field stimulation potential (which is Faraday-screened by the SS), or the AP elicited in SS membrane. Crucially, though, some TT electrically disconnected from the SS can be triggered to generate AP using optical stimulation (AP+; left panel in Figure 4B). The amplitude distribution of optically induced AP in SS and TT is depicted in Figure 4C. While AP amplitude in SS can be described by a unimodal distribution, TT exhibit a pronounced bimodal profile, where most TT are silent (AP-; right panel in Figure 4B) but a fraction of TT (AP+) are intrinsically excitable. Notably, the bimodal distribution of AP amplitudes in TT suggests that AP require activation of voltage-dependent channels with threshold activation (e.g. NaV1.5), rather than simply reflecting ChR2-induced inward currents. This is further confirmed in a patch clamp experiment on single cardiomyocytes, where the effect of ChR2-induced photocurrents on the membrane potential was evaluated in the presence or absence of Na^+^ and Ca^2+^ channel blockers (Figure S5).

**Figure 4:**
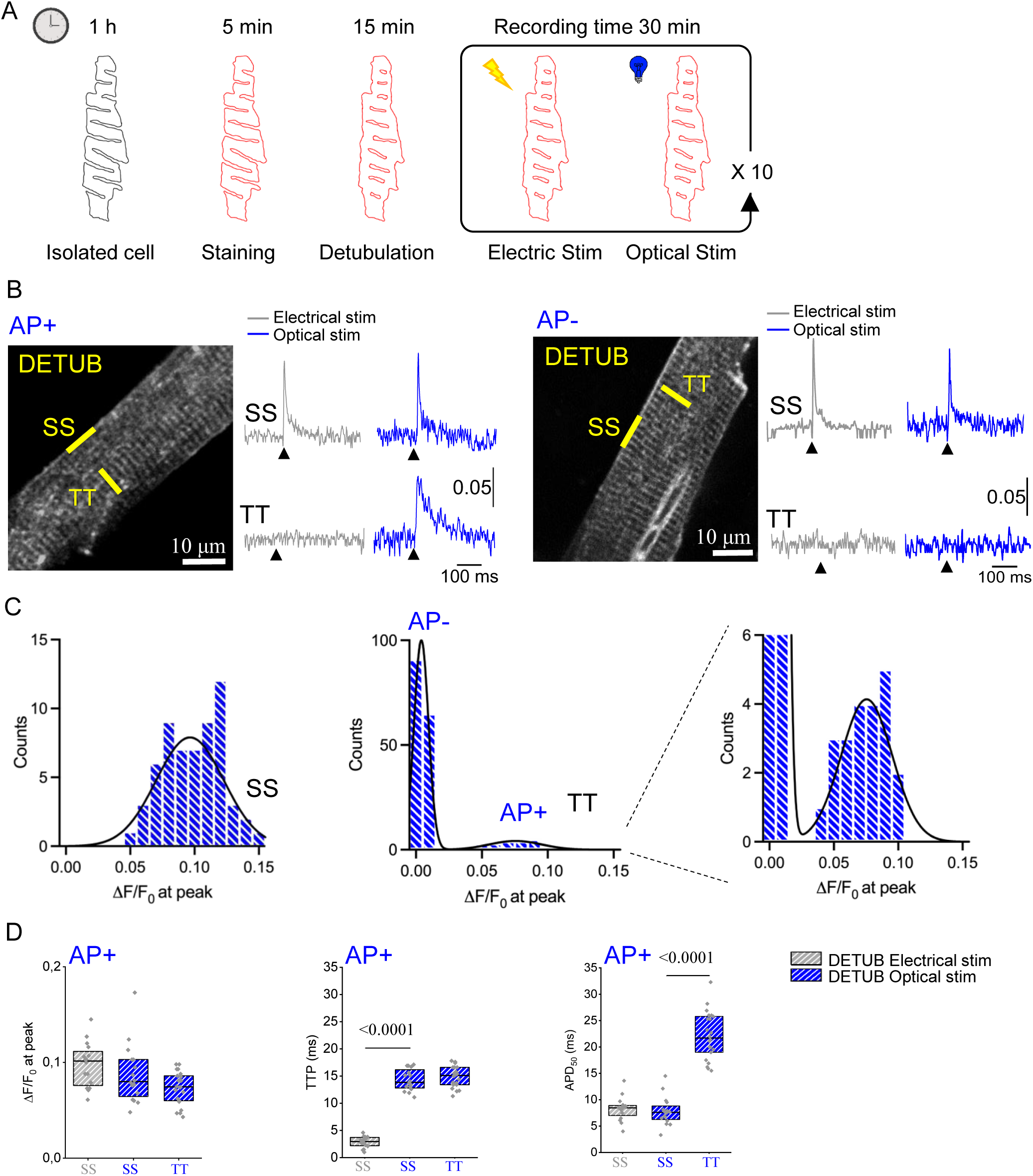
All-optical AP recording in TT und SS membrane sites of DETUB cardiomyocytes. (A) Representative scheme of DETUB, followed by electrical and optical AP stimulation. Isolated cardiomyocytes were stained using VSD (red), before acute detubulation. After DETUB, ten alternating cycles of electrical and optical stimulation were applied to elicit AP. (B) Two-photon fluorescence image of a DETUB ventricular myocyte constitutively expressing ChR2 and labelled with VSD. The yellow lines mark the probed membrane sites (sarcolemma: SS; T-tubules: TT). Scale bar 10 μm. On the side of the images, normalized fluorescence traces (ΔF/F_0_) recorded from the two scanned sites (average of 10 sequential trials) in DETUB cells. During electrical stimulation, an AP (elicited at 200 ms, black arrowheads) is clearly visible in SS, while detached TT do not generate AP (grey traces). Using light (blue traces), it is possible to trigger AP in SS, while TT shows double scenarios: the AP is elicited in detached TT (AP+), on the left; the AP is not triggered (AP-), on the right. (C) Distribution of AP amplitude (ΔF/F_0_ at peak) in both SS and TT. On the right a zoom in of TT AP amplitude to highlight the distribution of AP+. Data are from 68 SS and 178 TT (N = 8 animals, n = 68 cells). (D) AP amplitude (ΔF/F_0_ at peak), time-to-peak (TTP) and APD_50_, measured in DETUB cells upon electrical (grey) and optical (blue) stimulation. AP + and AP- is respectively refer to the membranes that are able or not to elicited an AP. Single data are reported as grey points and box plots with a range of 25^th^ to 75^th^ percentile are superimposed. Median values are represented by the line in box plots. Data are from 16 SS and 22 TT. ANOVA two ways with Bonferroni test was applied. Raw data are available at: https://doi.org/10.6084/m9.figshare.23742231.

In keeping with *in silico* predictions, isolated TT that are able to generate AP (AP+) show slower repolarization, compared to SS (increased AP duration at 50% repolarisation), unravelling intrinsic features of TT-generated AP (Figure 4D).

The fraction of TT that were optically excitable after osmotic detubulation (AP+) was only 10 - 15 % of all structurally disconnected TT (Figure 5A). We exclude that a low photo-current density would account for lack of TT excitability in the majority of TT, based on the homogenous sarcolemmal distribution of ChR2 measured in this transgenic mouse model (Figure 5B). However, a reduction of membrane excitability in sealed-off TT is not unexpected: luminal ion concentrations are no longer clamped to bulk extracellular levels and this can affect the local electrochemical gradient and thus channel/transporter functionality. To illustrate this, in Figure 5C we show a model prediction of how an increase in background Na^+^ permeability may lead to time-dependent variations of TT ion concentrations and resting membrane potential. For instance, if TT background Na^+^ permeability is increased 2.5 times, TT resting potential rises to :: -63 mV, so that the fraction of inactivated voltage-gated Na^+^ channel is increased from 0.13 to 0.96, and TT excitability is disabled (Moench & Lopatin, 2014). The simulated increase in TT background Na^+^ permeability has a phenomenological significance but do not reflect a unique molecular mechanism. Resting membrane potential variations of a similar extent can be related to variations of background permeability of K^+^ or other ions.

**Figure 5:**
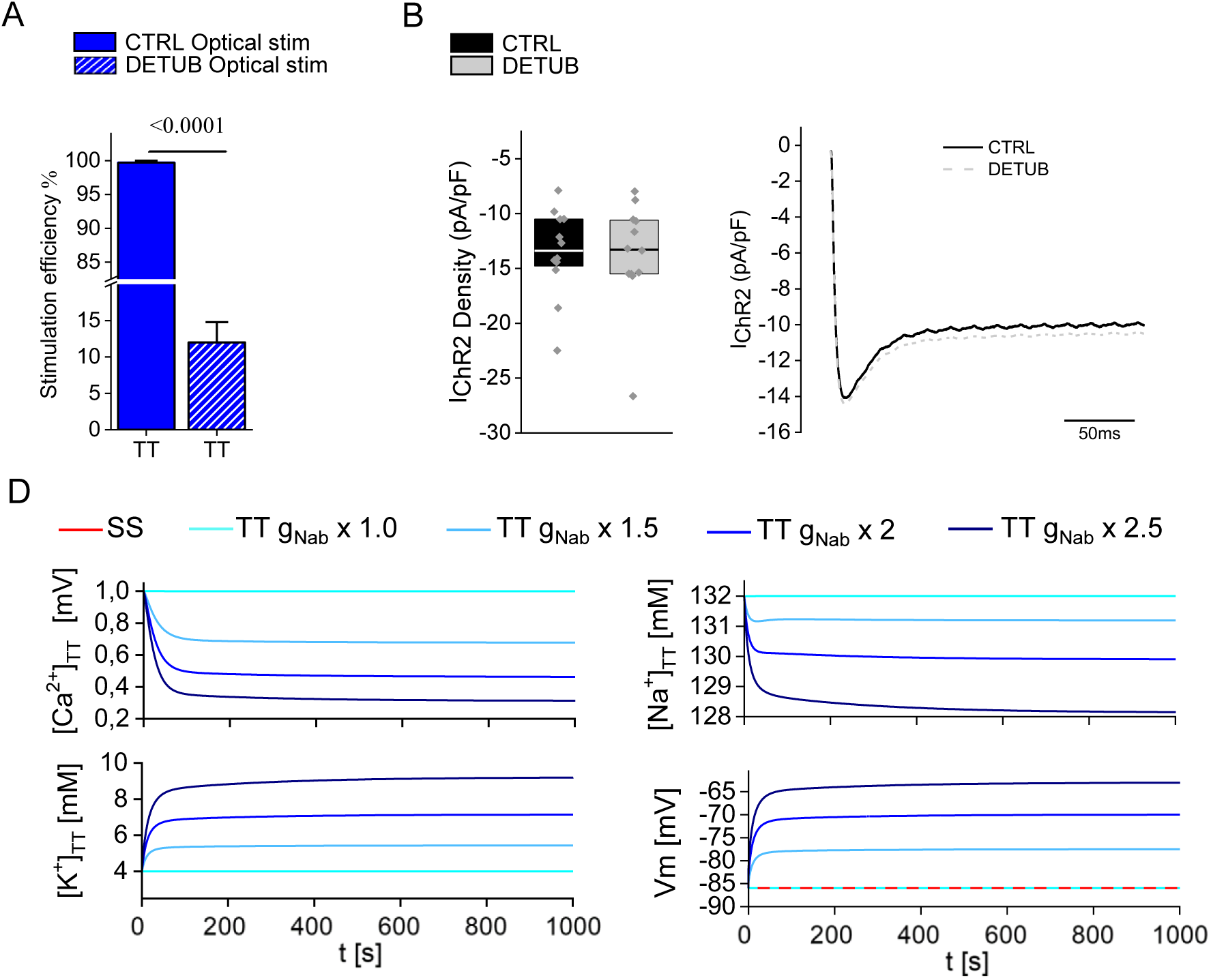
Loss of excitability of detached t-tubules. (A) Graph shows the percentage of TT in which optical stimulation can trigger AP CTRL and DETUB cells. Fisher exact test was applied. (B) Average value and representative traces of IChR2 density provoked using 0.4 mW of light intensity from CTRL (N = 3, n = 12) and DETUB (N = 3, n = 12), (C) Simulation of ion concentration changes and of membrane voltage over time in detached TT in control conditions (cyan line) and when background Na^+^ current (g_Nab_) was increased by 1.5× (light blue line), 2× (blue line), and 2.5× (dark blue line) at [Na^+^]_e_ =132 mM, [K^+^]_e_ = 4 mM and [Ca^2+^]_e_ = 1 mM. The model takes into account the experimentally used different levels of external ion concentrations during detubulation ([Na^+^]_e_ =133 mM, [K^+^]_e_ = 4.8 mM and [Ca^2+^]_e_ = 0.5 mM) and during the recordings ([Na^+^]_e_ =132 mM, [K^+^]_e_ = 4 mM and [Ca^2+^]_e_ = 1 mM). The transmembrane voltage at the SS is denoted by red line. Raw data are available at: https://doi.org/10.6084/m9.figshare.23742231.

## DISCUSSION

In this study, we exploit the utility of light to probe crucial electrophysiological properties of SS and TT membranes in living cardiomyocytes. Our approach combines the advantages of RAMP microscopy, in terms of spatio-temporal resolution for local AP recording, with optogenetics for precise spatio-temporally controlled stimulation of sub-cellular membrane domains. Applying this methodology to cardiomyocytes where TT were structurally disconnected from the SS membrane by DETUB, we confirm that TT membranes are electrically excitable – i.e. able to generate AP, locally and independently from the SS. We further identify dissimilarities in AP shape between the two membrane domains that are in keeping with differences in their Ca^2+^ current density: AP duration at 50% repolarisation is longer and AP-shape is more triangulated in TT than in SS. A complementary approach to verifying TT excitability might employ spatially-selective optogenetic stimulation of TT in an intact cardiomyocyte by two-photon excitation. However, due to the low density (and conductance) of ChR2 channels in the sarcolemma, it is very challenging to induce an AP upon femtoliter-range excitation volumes. Sophisticated beam-shaping systems (Papagiakoumou *et al*., 2010) would be needed to selectively spread the excitation light across an extensive enough region of the TT system.

We found that only a small fraction (10–15%) of seal-off TT are optically excitable. Although the full set of transporters and channels is present in the TT membrane of ventricular mouse cardiomyocytes (Thomas *et al*., 2003; Berry *et al*., 2007; Despa & Bers, 2007; Swift *et al*., 2007; Bossuyt *et al*., 2009; Gadeberg *et al*., 2017), in the limited lumen volume of vacuolated TT, ion concentrations can largely fluctuate both during cell activity and long quiescence periods. Indeed, accumulation of K^+^ and depletion of Ca^2+^ have been predicted and experimentally measured (Pasek *et al*., 2006; Moench & Lopatin, 2014). In our in silico investigation, ion fluctuations were phenomenologically introduced by increasing TT Na^+^ background permeability. Indeed, in vacuolated TT, local diastolic Na^+^ concentrations in the intra- and extracellular space are continuously reset by the balance between passive Na^+^ entry (through voltage- or stretch-activated Na^+^ channels and forward-mode NCX) versus active Na^+^ efflux (mainly through NKA). The behavior of these channels and transporters in vacuolated TT is particularly difficult to predict and potentially complicated by the presence of neuronal isoforms of voltage-gated Na^+^ channels (i.e. Nav1.1, Nav1.3, and Nav1.6 are enriched in T-tubules and have a lower activation threshold and inactivate more rapidly). We computationally observed that, in these vacuolated TT, a minimal unbalance toward an increased background Na^+^ influx (versus efflux) can produce a resting membrane potential depolarization of 10-15 mV, sufficient to inactivate the large quota of voltage-gated Na^+^ channels and thus to compromise membrane excitability. Similar variations of vacuolated TT membrane resting potential can be predicted if background K^+^ fluxes are unbalanced as a consequence of K^+^ accumulation in the TT lumen (and consequent increase of K^+^ equilibrium potential), as well as, also in this case, variations of NKA activity.

TT membrane excitability had previously been reported for skinned fibres of vertebrate skeletal muscle (Posterino *et al*., 2000; Ortenblad & Stephenson, 2003; Macdonald & Stephenson, 2004; Pedersen *et al*., 2005). In those experiments, the SS of muscle fibres was mechanically removed by rolling back the surface sarcolemma with forceps, forming a sort of ‘cuff’, while part of the TT system sealed off. In our report on isolated living cardiomyocytes, electrical uncoupling between SS and TT was achieved using transient osmotic shock that physically detaches TT membranes from the cell surface, while in principle maintaining overall cell integrity, with intact SS and TT system components that are still operational, and a near-native cellular environment.

In contrast to skeletal muscle, cardiomyocyte activation *in vivo* occurs via electrotonic interaction with neighbouring cells, AP duration is longer (by one to two orders of magnitude, depending on species), and ECC involves trans-sarcolemmal calcium fluxes, making cells highly responsive to changes in local ion concentrations and regulatory influences on ion flux pathways.

Our study shows that TT do not merely represent a “current sink” compartment that is driven by the SS “source”, but rather that TT are part of an overall excitable membrane system in which SS and TT affect each other. Several considerations of the physiological and patho-physiological consequences of the present findings can be made.

The capacity of TT to generate AP may protect cells from the effects of spatio-temporal heterogeneities of membrane depolarisation that may occur within the TT network, in particular when the TT system is structurally remodelled, such as in disease. In healthy cardiomyocytes, the characteristic length constant of electrotonic conduction along TT membranes is one order of magnitude larger than cell radius (Scardigli *et al*., 2017; Kong *et al*., 2018), but the utility of this length constant as a measure to predict spatial voltage uniformity within the TT network is questionable in supra-threshold voltage ranges (because membrane resistance markedly varies during the AP). The scenario becomes even more complex in in atrial cardiomyocytes, where the tubular system is less developed and mainly composed of axial elements (Richards *et al*., 2011; Brandenburg *et al*., 2016; Brandenburg *et al*., 2018). This complexity is also evident in disease settings, where the spatial density of the TT system is altered, and its structural integrity is compromised (Ibrahim *et al*., 2011). While in healthy ventricular cardiomyocytes no significant voltage gradient is expected, in pathologically remodelled tubular system the resistivity of the extracellular space may significantly increase and tubular elements can be poorly coupled, or even electrically uncoupled from the SS.

The pathophysiological relevance of TT excitability remains to be elucidated. Interestingly, behaviour similar to what has been observed here has been reported previously in pathological conditions (Crocini *et al*., 2014; Crocini *et al*., 2016b). Thus, in heart failure, when the structural integrity of the TT system is compromised, some TT elements are apparently electrically silent and/or not able to generate AP in synchrony with the SS. However, a number of these TT elements show local spontaneous electrical activity, indicative of TT membrane excitability (Sacconi *et al*., 2012). These spontaneous TT depolarizations do not propagate to the SS, and they may trigger large local SR Ca^2+^ releases (voltage-associated Ca^2+^ sparks, V-sparks) (Crocini *et al*., 2014). Future investigations will focus on exploring the relevance of TT excitability in cardiomyocytes whose ECC homogeneity has been challenged as a consequence of a pathologically remodelled TT system, employing the experimental approach described here to dissect TT and SS excitability and AP features. At this point, however, we can unequivocally confirm that TT membranes in intact cardiomyocytes are electrically excitable.

## ACKNOWLEDGEMENTS

This work was supported by the European Union’s Horizon 2020 research and innovation program under grant agreement no 871124 Laserlab-Europe. P Kohl and L Sacconi are members of the German Research Foundation’s Collaborative Research Centre SFB1425 (DFG #422681845). The work of M Pásek was carried out with the Institutional Support RVO: 61388998.

