## Supplementary material for "Optogenetic confirmation of transverse-tubular membrane excitability in intact cardiac myocytes"

### 1. Reformulation of the model of rat to mouse ventricular myocyte

A biophysically detailed computer model of a mouse ventricular cardiomyocyte, incorporating both surface sarcolemma (SS) and t-tubular membrane (TT) was used (Figure S2). The TT network was modelled as a single compartment, separated from the surface membrane by the mean resistance of the TT system, and connected to the bulk extracellular solution.

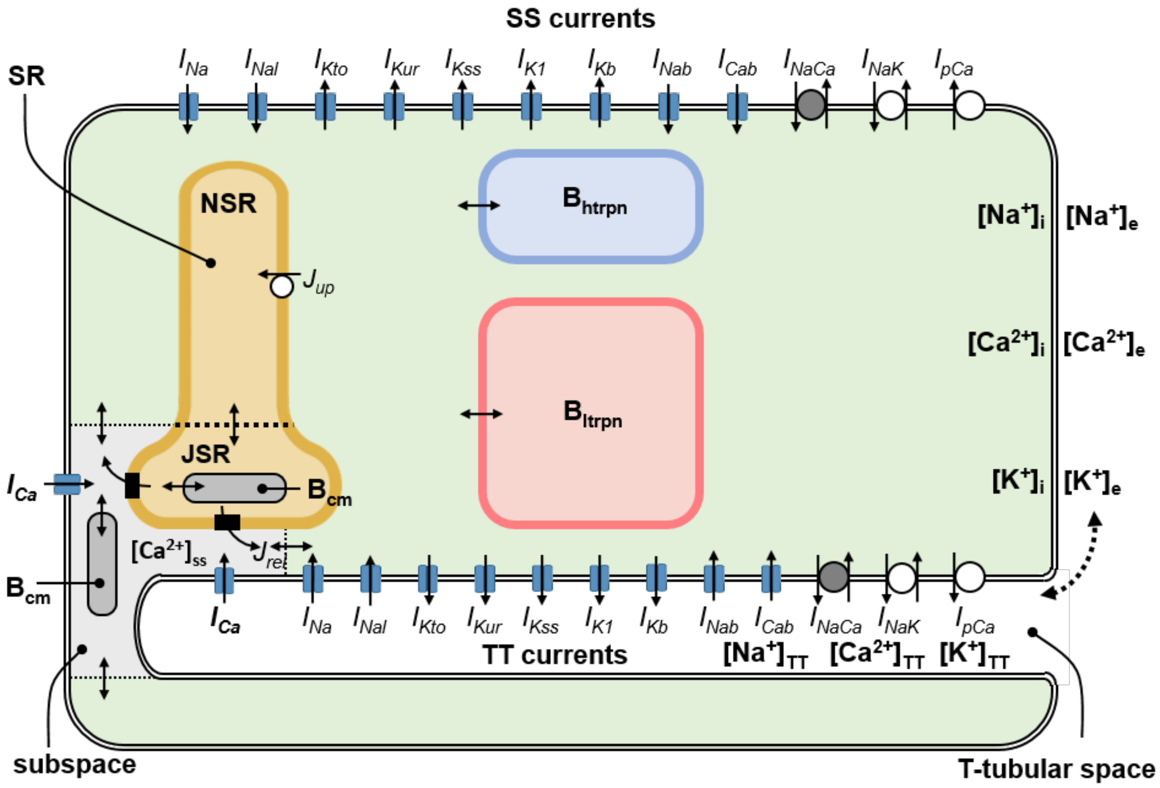


**Figure S1: Schematic diagram of the mouse ventricular myocyte model.** The description of electrical activity of SS and TT comprises formulation of the following ion currents: fast $Na^+$ current ($I_{Na}$), late $Na^+$ current ($I_{NaL}$), L-type $Ca^{2+}$ current ($I_{Ca}$), time-dependent transient outward $K^+$ current ($I_{Kto}$), ultrarapidly activating delayed rectifier $K^+$ current ($I_{Kur}$), noninactivating steady-state $K^+$ current ($I_{Kss}$), time-independent inward rectifier $K^+$ current ($I_{K1}$), background $Na^+$ current ($I_{Nab}$), background $Ca^{2+}$ current ($I_{Cab}$), background $K^+$ current ($I_{Kb}$), $Na^+$- $Ca^{2+}$ exchange current ($I_{NaCa}$), $Na^+$- $K^+$ ATPase pump current ($I_{NaK}$), sarcolemmal $Ca^{2+}$ pump current ($I_{pCa}$). The intracellular space contains the subspace, the network and junctional compartments of sarcoplasmic reticulum (NSR, JSR) and the $Ca^{2+}$ buffers calmodulin ($B_{cm}$), troponin ($B_{htrpn}$, $B_{ltrpn}$) and calsequestrin ($B_{cs}$). The small filled rectangles in JSR membrane represent ryanodine receptors. The small bi-directional arrows denote $Ca^{2+}$ diffusion and the thick dashed arrow represents ionic diffusion between the TT and the external bulk spaces.

#### *2.1 Geometric parameters of cell and of cellular compartments*

The total volume of model cell ($V_c$) was set to 10 pL. The fractional volumes of cytosolic space and dyadic space, as well as of network and junctional (release) compartments of the sarcoplasmic reticulum (NSR, JSR) are as in original rat model (Pasek *et al.*, 2006): 0.585, $75\times10^{-6}$, 0.0315 and 0.0035, respectively. The fractional volume of TT was set to 3.2% (Clark *et al.*, 2001).

*2.2 Membrane Capacitance*

According to our experimental data, the mean values of membrane capacitance ($C$) in intact ($C_{CTRL}$) and detubulated ($C_{DETUB}$) cells were 193 and 126 pF, respectively, and the mean fraction of TT resisting detubulation ($TT_{res}$) was 0.3. Assuming that the specific membrane capacitance is comparable in both membrane domains, the following equations allow one to calculate the membrane capacitance in TT and SS ($C_{TT}$ and $C_{SS}$).

$$C_{CTRL} = C_{SS} + C_{TT} \quad (1)$$

$$C_{DETUB} = C_{SS} + TT_{res} \cdot C_{TT} \quad (2)$$

$$C_{CTRL} = 193\, pF;\ C_{DETUB} = 126\, pF;\ TT_{res} = 0.3\ \Rightarrow C_{SS} = 97.3\, pF;\ C_{TT} = 97.3\, pF \quad (3)$$

The conclusion that about half of the total sarcolemmal surface is in TT in mouse ventricular cardiomyocytes is consistent with a previous optical study by Forbes *et al.* (Forbes *et al.*, 1984). Applying this data to a model with a total membrane area of 10,000 μm$^2$ and a specific membrane capacitance of 1 μF/cm$^2$, the modelled TT and SS have an identical capacitance of 50 pF. Please note that the behaviour of the model is independent of the size of the cell considering that everything is bound by mutual ratios.

*2.3 Membrane currents*

To simulate the behaviour of mouse ventricular septal cardiomyocyte (Anumonwo *et al.*, 2001), the following modifications of membrane transport systems were implemented.

(i) The conductance of $I_{Na}$-channels ($g_{Na}$) was increased from 10 to 13.3 mS/cm$^2$ to adjust maximum upstroke velocity to ~150 V/s as observed experimentally (Anumonwo *et al.*, 2001). Additionally, the membrane $Na^+$ transport was supplemented by a formulation of late $Na^+$ current ($I_{NaL}$), adopted from the mouse ventricular myocyte model of Morotti *et al.* (Morotti *et al.*, 2014). The corresponding conductance ($g_{NaL}$) was set to 6.5 μS/cm$^2$.

(ii) The permeability of $I_{Ca}$-channels ($P_{Ca}$) was decreased from $2.1\times10^{-4}$ to $0.7\times10^{-4}$ cm/s and the time constant of voltage dependent activation of $I_{Ca}$ ( $\tau_{Ca,VDA}$) was adjusted to reduce the rate of $I_{Ca}$ deactivation in later phases of AP repolarisation (negative to -20 mV), allowing the model AP duration at 75% repolarisation to approach the value of 20.9 ±1.3 ms, measured in mouse ventricular cardiomyocytes at 1 Hz stimulation (Guo *et al.*, 1999).

(iii) The conductance of $I_{Kto}$-channels ($g_{Kto}$) was decreased from 0.35 to 0.146 mS/cm$^2$ and gating characteristics of the channels were reformulated to mimic the parameters of early AP repolarization ($APD_{25}$ = 4 ± 0.2 ms and $APD_{50}$= 8.5 ± 0.3), measured at 1 Hz (Guo *et al.*, 1999).

(iv) The formulations of $I_{Ks}$-channels (slow component of the delayed rectified $K^+$ current) and of the newly added $I_{Kur}$-channels (ultra-rapid component of the delayed rectified $K^+$ current) were adopted from the model of mouse ventricular myocytes developed by Pandit *et al.* (Pandit, 2003) that is available in the CellML model repository (https://models.cellml.org/cellml). The related conductances $g_{Kss}$ and $g_{Kur}$ were set to 0.075 and 0.085 mS/cm$^2$, respectively.

(v) Other changes to the original rat model include: decrease of $g_{K1}$ from 0.24 to 0.159 mS/cm$^2$, omission of hyperpolarization-activated current ($I_f$), increase of $g_{Nab}$ (background $Na^+$ conductance) from 0.8 to 1.2 μS/cm$^2$, decrease of $g_{Cab}$ (background $Ca^{2+}$ conductance) from 0.648 to 0.367 μS/cm$^2$ (Anumonwo *et al.*, 2001), increase of $I_{NaK,max}$ (maximal density of $Na^+$-$K^+$ ATPase current) from 1 to 1.4 μA/cm$^2$ and of $Na^+$-half-saturation constant for $I_{NaK}$ from 10 to 21 mM (Anumonwo *et al.*, 2001), as well as decrease of $I_{pCa,max}$ (maximal density of sarcolemmal $Ca^{2+}$-pump current) from 0.85 to 0.21 μA/cm² (summarised in Table T1).

Consistent with the absence of changes in $I_{Na}$, $I_{K,}$ and membrane resting potential after detubulation as well as with excess of $I_{Ca}$ at the TT membrane (see Figure S1), we simplified the distribution of ion transporters by setting all fractions to 0.5 (uniform distribution between the TT and SS membranes), with the exception of L-type $Ca^{2+}$ current ($f_{Ca}$).

$f_{Ca}$ was estimated on the basis of measurements of membrane capacitance and corresponding current densities in intact and detubulated cardiomyocytes ($C_{CTRL}$, $C_{DETUB}$, $I_{x,CTRL}$, $I_{x,DETUB}$) and relations derived in Pásek at al. (Pasek *et al.*, 2017; Pasek *et al.*, 2021):

$$I_{x,ratio} = \frac{I_{x,CTRL}C_{CTRL} - I_{x,DETUB}C_{DETUB}}{I_{x,CTRL}C_{CTRL}TT_{res} - I_{x,DETUB}C_{DETUB}} \cdot \frac{C_{CTRL}TT_{res} - C_{DETUB}}{C_{CTRL} - C_{DETUB}} \quad (4)$$

$$f_x = \frac{I_{x,ratio}}{C_{SS}/C_{TT} + I_{x,ratio}} \quad (5)$$

where $I_{x,ratio}$ represents the ratio between current densities in the TT and SS membranes ($I_{x,TT}/I_{x,SS}$), $TT_{res}$ stands for the fraction of TT resisting detubulation, and $C_{TT}$ and $C_{SS}$ represent the membrane capacitance of the respective membrane domains.

Assuming that $C_{TT} \approx C_{SS}$, and taking into account the mean values of $C_{CTRL}$ and $C_{DETUB}$ in a subpopulation of cells used for our $I_{Ca}$ recording ($C_{CTRL}$ = 162 pF, = 97 pF), we conclude that the $TT_{res}$ was ~ 0.2 in the corresponding cells (see equations 1 and 2). Using the mean values of $I_{Ca,CTRL}$ and $I_{Ca,DETUB}$ (6.5 and 3.26 pA/pF, respectively) and inserting these data into equations 4 and 5 gives $I_{Ca,ratio}$ of 6.9 and $f_{Ca}$ of 0.87, which corresponds well with the previous estimates done by Gadeberg *et al.* (Gadeberg *et al.*, 2017) suggesting predominant localisation of $I_{Ca}$-channels in TT of mouse ventricular cardiomyocytes.
The electrical properties of all ion transport systems used in the model are summarized in Table T1.

**Table T1:** Electrical properties of ion transport systems in the model of mouse ventricular myocyte

| *parameter* | *value* | $f_x$ | *parameter* | *value* | $f_x$ |
|---|---|---|---|---|---|
| $g_{Na}$ | 13.3 mS/cm² | 0.5 | $g_{Nab}$ | 1.2 μS/cm² | 0.5 |
| $g_{NaL}$ | 6.5 μS/cm² | 0.5 | $g_{Cab}$ | 0.367 μS/cm² | 0.5 |
| $P_{Ca}$ | 0.7×10⁻⁴ cm/s | 0.87 | $g_{Kb}$ | 1.380 μS/cm² | 0.5 |
| $g_{Kto}$ | 0.146 mS/cm² | 0.5 | $k_{NaCa}$ | 0.18 nA/cm²/ mM⁴ | 0.5 |
| $g_{Kss}$ | 0.075 mS/cm² | 0.5 | $I_{NaK,max}$ | 1.4 μA/cm² | 0.5 |
| $g_{Kur}$ | 0.085 mS/cm² | 0.5 | $I_{pCa,max}$ | 0.21 μA/cm² | 0.5 |
| $g_{K1}$ | 0.159 mS/cm² | 0.5 | | | |

Parameters of individual transporters (conductance ($g$), permeability ($P$), maximum current ($I_{XX,max}$) or scaling factor ($k_{NaCa}$)). $f_{x,t}$ represents the TT fraction of the ion transporter underlying current $I_X$.

### *2.4 Intracellular $Ca^{2+}$- handling*

To simulate the increase of the systolic and diastolic maxima of intracellular $Ca^{2+}$ transients with stimulation frequency observed in mouse ventricular cardiomyocytes by Ito *et al.* (Ito *et al.*, 2000), the forward rate parameter of SR $Ca^{2+}$ ATPase was decreased from 0.4 to 0.2 mM/s and the slope factor controlling the $[Na^+]_i$ dependence of $Na^+/K^+$ pump was increased from 1.5 to 2.15.

*2.5 Rate of ion exchange between TT and external space*

The time constant of $K^+$ exchange between the TT and external space ($\tau_{K,te}$) was set to 50 ms to comply with the average rate of tail current decay observed at the end of depolarising steps from -80 mV to the voltages between -30 and 40 mV (Clark *et al.*, 2001). $\tau_{Na,te}$ was set to the same value as $\tau_{K,te}$ and $\tau_{Ca,te}$ was set to 167 ms to respect the ratio $\tau_{Ca,te} / \tau_{K,te}$ in the original rat model (Pasek *et al.*, 2006).

*2.6 Basic behaviour of the model*

The basic behaviour of the mouse ventricular cell model is illustrated in Fig. S3. It shows superimposed action potential (AP) shapes, principal membrane currents, and ion concentrations in intracellular and extracellular compartments during 1, 3 and 5 Hz steady-state stimulation (after 600 s of stabilisation).

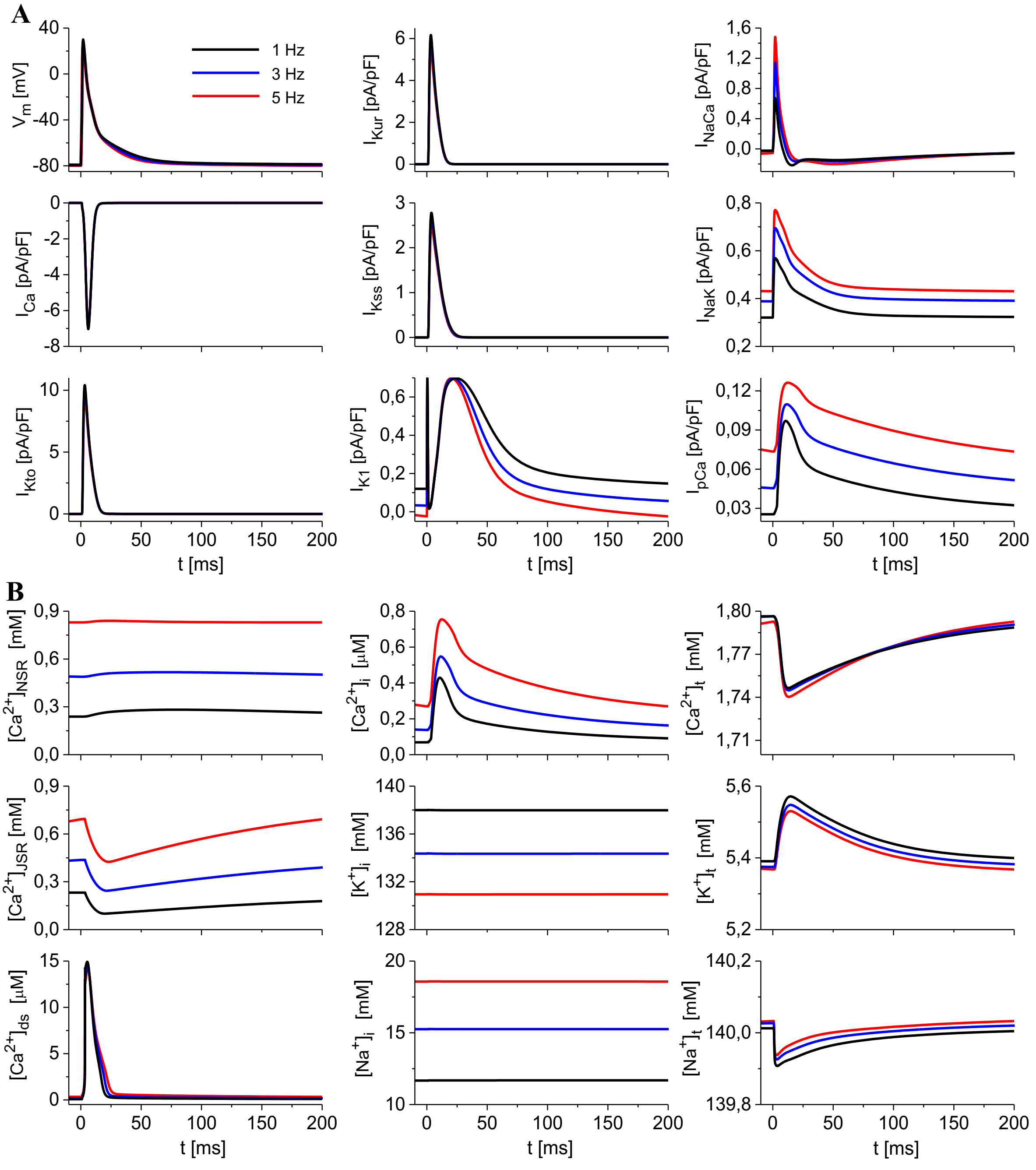

**Figure S2:** Behaviour of the model of a mouse ventricular cardiomyocyte at 1, 3 and 5 Hz steady-state stimulation. A) Membrane voltage ($V_m$) and ion currents ($I_{Na}$, $I_{Ca}$, $I_{Kto}$, $I_{Kur}$, $I_{Kss}$, $I_{K1}$, $I_{NaK}$, $I_{NaCa}$, $I_{pCa}$,). B) Ion concentration changes in network and junctional (release) compartments of the SR ($[Ca^{2+}]_{NSR}$, $[Ca^{2+}]_{JSR}$), in dyadic space and bulk cytoplasm ($[Ca^{2+}]_{ds}$, $[Ca^{2+}]_i$, $[K^+]_i$, $[Na^+]_i$) and in TT ($[Ca^{2+}]_t$, $[K^+]_t$, $[Na^+]_t$). Extracellular ion concentrations were set to physiological values: $[Na^+]_e$ = 140 mM, $[K^+]_e$ = 5.4 mM and $[Ca^{2+}]_e$ = 1.8 mM.

Similarly to the model by Bondarenko *et al.* (Bondarenko *et al.*, 2004), the characteristics of AP reproduced by our model ($APD_{25}$ = 4 ms, $APD_{50}$ = 8 ms and $APD_{75}$ = 16 ms at 1 Hz) agree well with experimentally measured values (Guo *et al.*, 1999) and change only slightly with stimulation frequency (Guo *et al.*, 2000). The model reconstruction of experimental data of Ito *et al.* (Ito *et al.*, 2000), showing a frequency-dependent increase in the systolic and diastolic maxima of intracellular $Ca^{2+}$ transients, is illustrated in Fig. S4.

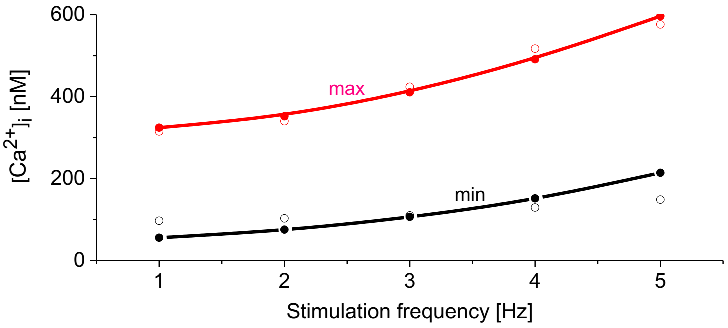


**Figure S3:** Model reconstruction of the frequency dependence of intracellular $Ca^{2+}$ transient amplitudes in mouse ventricular myocytes published by Ito *et al.* (Ito *et al.*, 2000). Black and red empty circles represent minimal (diastolic) and maximal (systolic) values of $Ca^{2+}$ transients in experiments, respectively. Black and red full circles represent corresponding data from the model. Extracellular ion concentrations in experiment and model: $[Na^+]_e$ = 137 mM, $[K^+]_e$ = 3.7 mM and $[Ca^{2+}]_e$ = 1.5 mM.

### 2. Adeno-associated virus-based gene transfer of CheRiff

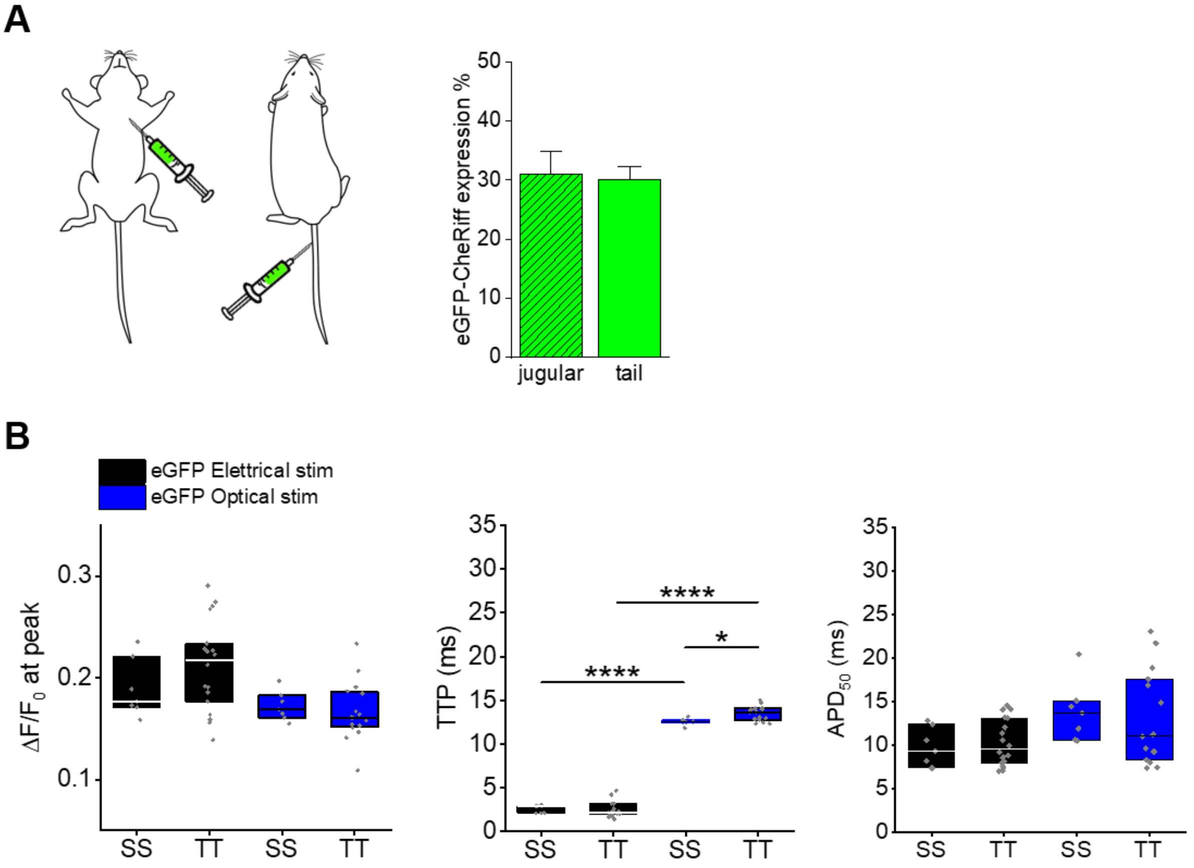


**Figure S4: eGFP-Cheriff characterization.** (a) Sketches show two types of viral injections: left into the jugular and right into the tail vein. Columns show the comparison of eGFP expression percentage in cardiomyocytes isolated from mice injected with the two approaches. No significant difference was observed (P = 0.8225). Student's t-test; n =146 cells with jugular and n = 404 with tail vein injections; cells were isolated from N = 2 mice for each approach). (b) Columns showing AP amplitudes ($\Delta F/F_0$ at peak), time-to-peak (TTP), and AP duration at 50% repolarization ($APD_{50}$) mean values measured in eGFP-Cheriff cells using electrical field stimulation (black) as compared to optical stimulation (blue). Single cell data are reported as grey points and box plots with a range of $25^{th}$ to $75^{th}$ percentile are superimposed. Median values are represented by the line in box plots. Data from 7 SS and from 18 TT recordings (n = 7, N = 3). P values were calculated with two-way ANOVA with Bonferroni test. ( *p < 0.05, ****p < 0.0001). N indicates the number of animals and n the number of cells in each data series.

3. **Membrane potential variation upon ChR2 stimulation.**

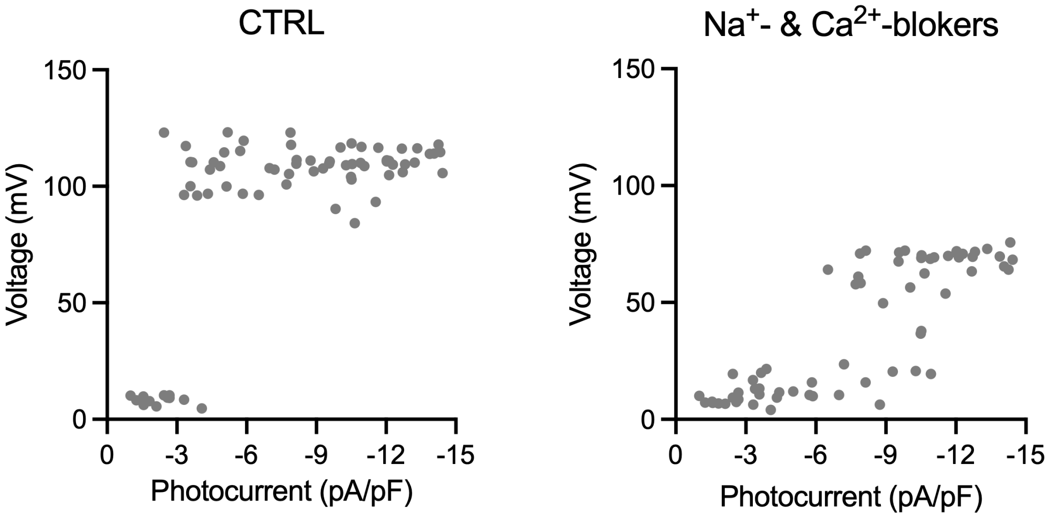


**Figure S5:** Figure S5: Characterization of membrane potential variation versus light-induced ChR2 photocurrent in CTRL cells in the absence and after treatment with TTX (at 30 µM) and lacidipine (10 µM). Data from 19 cells (4 mice).
